# Long-term community change through multiple rapid transitions in a desert rodent community

**DOI:** 10.1101/163931

**Authors:** Erica M. Christensen, David J. Harris, S. K. Morgan Ernest

## Abstract

While studies increasingly document long-term change in community composition, whether long-term change occurs gradually or via rapid reorganization events remains unclear. We used Latent Dirichlet Allocation (LDA) and a change-point model to examine the long-term dynamics of a desert rodent community undergoing compositional change over a 38-year span. Our approach detected three rapid reorganization events, where changes in the relative abundances of dominant and rare species occurred, and a separate period of increased variance in the structure of the community. These events coincided with time periods—possibly related to climate events—where the total abundance of rodents was extremely low. There are a variety of processes that could link low abundance events with a higher probability of rapid ecological transitions, including higher importance of stochastic processes (i.e., competitive interactions or priority effects) and the removal of structuring effects of competitive dominants or incumbent species. Continued study of the dynamics of community change will provide important information not only on the processes structuring communities, but will also provide guidance for forecasting how communities will undergo change in the future.

## Introduction

As humans alter the template of nature by increasing temperature, changing nutrient distributions, and altering land cover (Walther 2010), the composition of species living in these places also changes. Compositional changes occur both directly as each species responds to the environment in the context of its own needs or preferences, and indirectly through changes in the competitive landscape and other species interactions. Depending on the mechanisms, community change can be gradual or rapid. Gradual change can occur through stochastic turnover events, or niche-based turnover as species’ ability to thrive gradually improves or degrades as the environment changes (e.g. Tingley et al. 2009). Rapid changes in communities can emerge as ecosystems respond to intrinsic or extrinsic drivers (Williams et al. 2011). With intrinsically-driven rapid change, sometimes referred to as regime shifts, gradual changes in the environment eventually push the ecosystem past a threshold, triggering rapid shifts to an alternate stable state (Scheffer and Carpenter 2003). Extrinsically-driven rapid changes can occur either from niche-based tracking as the environment rapidly shifts from one state to another (Beaugrand 2004), or via extreme events, which cause cascading changes in species populations that alter how the community recovers post-disturbance (Smith 2011).

While there are growing numbers of studies documenting that communities are changing, we know little about the dynamics of how this change is occurring. Meta-analyses compiling data from many long-term studies have focused on the basic question of whether communities are changing (La Sorte and Boecklen 2005, Dornelas et al. 2014), with little or no focus on the pattern of that change (but see Bagchi et al. 2017). A growing number of studies document the occurrence of rapid ecological transitions (Beaugrand 2004, Thibault and Brown 2008), but whether these events are rare or a common mechanism by which long-term change occurs is unclear. This may be because studies focused on rapid ecological transitions are asking different questions than meta-analyses examining long-term community change. Rapid ecological transitions are unpredictable and abrupt, thus published studies typically focus on relatively short time-scales and include only the dynamics immediately before and after the specific event of interest (e.g. Thibault and Brown 2008). In contrast, meta-analyses of community change focus on the trends occurring over decades, and may intentionally average out or avoid periods of rapid transition. Because studies of rapid ecological transitions and meta-analyses are focused on different patterns at different time scales, it is unknown whether rapid ecological transitions are truly rare events. Answering this question requires high frequency, long-term monitoring data and methods that are able to detect these dynamics in ecological time series.

Here, we examine community change through time in a desert rodent community. Surveyed monthly from 1977 to 2015, this rodent community has undergone significant turnover. Both rapid and gradual reorganization have been invoked to explain these changes in rodent composition. Studies focused on rodent species turnover have interpreted dynamics as indicative of gradual long-term response to habitat shifts from open arid grassland to shrubland (Ernest et al. 2008). Other studies, focused on the impacts of specific climate events on the rodent community, have proposed that these events triggered rapid shifts in community composition (Valone et al. 1995, Thibault and Brown 2008). Therefore this is an ideal data set for assessing how to reconcile short-term events with long-term community change at the multi-decade scale.

## Methods

To examine the dynamics of this community, we used 38 years of monthly rodent data from the Portal Project, a long-term study located on 20 hectares of Chihuahuan Desert near the town of Portal, Arizona, USA (Ernest et al. 2016). This site has undergone considerable habitat change: a 3-fold increase in woody vegetation between 1977 and the mid-1990s transitioned this site from an open desertified grassland with widely scattered woody shrubs to a desert shrubland (Brown et al. 1997). Small mammal data is collected at this site on 24 permanent 50m by 50m plots, sampled at monthly intervals with no major changes in methodology since 1977. Here, we pool data from the 8 unmanipulated (“control”) plots, which allow unrestricted access to all species from the regional pool, to provide one site-level estimate of the natural dynamics of the rodent community from 1977-2015. Data for our analysis consisted of a table of counts for each species for each month of the time series. This amounted to 436 time steps and 21 species.

The twenty-one rodent species caught at the site consist primarily of granivorous rodents, with some insectivorous or folivorous species. The body size range of this community spans from 6 grams to nearly 200 grams. This community has experienced considerable turnover in species composition through time, with only two species captured consistently at almost every survey since 1977. Further information on the site and protocol can be found in Brown (1998) and Ernest et al. (2016). The latter paper also contains the data used in this analysis.

To quantify the dynamics of species composition, we used a 2-step approach. First, we reduced the dimensionality of composition data (i.e. condensed the species-level information on presence/absence and abundances) using a method from machine learning (Latent Dirichlet Allocation; see Blei et al. 2003). We then examined the dynamics of this simplified composition time series using a change-point model (Western and Kleykamp 2004). We demonstrate the 2-step approach with a set of simulated data in Appendix S1.

### Identifying species associations using Latent Dirichlet Allocation

Latent Dirichlet Allocation (LDA) was developed as an alternative to cluster analysis for summarizing documents based on the words they contain. This machine learning approach was recently introduced to ecology as a way to quantify changes in species composition across gradients (see Valle et al. 2014). LDA takes the words in a document and identifies “topics” – collections of words that tend to be found together in specific proportions (Blei et al. 2003). Whereas clustering approaches assign each sample (“document”) to exactly one cluster, LDA attempts to infer the relative contributions of all topics. If these contributions change gradually over a series of documents, then these changes in document composition can be tracked from sample to sample. When applied to ecological data, this technique can summarize community observations (“documents”) based on the species (“words”) they contain. Analogous to collections of words forming topics in document analysis, LDA identifies groups of species that tend to be found together in specific proportions. Rather than looking at raw species composition in a collection of community observations (for example over a gradient in space or time), LDA is able to: 1) reduce species composition at each sampling point to its community-type composition, and 2) describe each community-type in terms of the species it contains, which may reveal important information about underlying species associations.

Fitting an LDA model involves simultaneously determining two sets of numbers: one defining the community-types, and one describing the observed species assemblages in terms of those types. Due to the complex relationship between the way each community-type is defined and its influence on individual assemblages, exact inference is not possible in this model. We used Blei et al.’s variational approximation, which simplifies this relationship. This approximation allowed us to use Blei et al.’s (2003) iterative “variational expectation-maximization” procedure for jointly optimizing both sets of parameters, as implemented in the `topicmodels` package (version 0.2-7) (Hornik and Grün 2011) for R 3.3.2 (R Core Team 2016). Like many clustering and ordination methods, LDA requires the number of community-types to be specified as model input (it does not determine the number of community-types supported by the data). We used AIC to select the appropriate number of community-types needed to parsimoniously describe the data (Appendix S2: Fig. S1). We calculated this value using the log-likelihood returned by the ‘LDA’ function in the ‘topicmodels’ package, penalized according to the number of free parameters. This approximation likely overestimates the number of parameters and therefore could underestimate the number of topics, but our qualitative time series results are robust to including more topics than the number selected. See Valle et al. 2014 for a thorough description of LDA in an ecological context, and comparison of LDA to traditional clustering techniques.

### Quantifying when change occurs using a change-point model

LDA is a discovery tool which simplifies multivariate species composition to better visualize community dynamics. However, it does not tell us if (or when) a change in community structure has occurred. We fit a change-point model (Western and Kleykamp 2004) to identify abrupt transitions in the time series of community-types generated by the LDA model. Change-point models break up a time series into intervals, with a different set of parameters to describe the time series during each interval. Between each pair of change-points, we modeled each community-type’s prevalence as a sinusoid with a period of one year, to control for seasonal fluctuations. The community-type proportions in each interval were modeled using a separate multinomial generalized linear model fit with the nnet package (Venables and Ripley 2002). Because we were modeling proportions rather than counts, this model gave us a *quasi*-likelihood for each interval, rather than a conventional likelihood (McCullagh and Nelder 1989). Since our quasi-likelihoods did not inflate the variance, however, they can be interpreted on the same scale as a conventional likelihood for purposes of model comparison (Anderson et al. 1994). The product of interval-level quasi-likelihoods yields the quasi-likelihood for the full data set; this value will be largest when the change-points break the time series into relatively stable intervals that can be explained well by the generalized linear model.

The number of possible change-point locations was too large to evaluate all the possibilities exhaustively, so we used Markov chain Monte Carlo (MCMC) to collect a representative sample of change-point locations that are consistent with the data. Initial experiments with Metropolis sampling showed poor mixing, so we implemented a parallel tempering sampler (also called Metropolis-coupled MCMC and replica-exchange MCMC) to facilitate movement of the Markov chain between modes via exchanges with auxiliary Markov chains that rapidly explore the space of possible change-points (Earl and Deem 2005). We fit change-point models with up to five change-points for the rodent data, and evaluated model performance by comparing the average log-quasi-likelihood to the number of model parameters (Gelman et al. 2014).

Our change-point model only detects changes in the mean value or seasonal amplitude of the community-types’ time series. Real dynamics in nature are likely a complicated combination of changes in mean, value, slope, and seasonal amplitude, making realistic models difficult to develop and interpret. Our change-point model does not explicitly look for changes in slope (linear trends), but the rate of change can be inferred by examining the distributions surrounding the change-points identified by the model. Figure S2 demonstrates that a rapid change in mean value of two community-types results in a narrow distribution. As the transition time lengthens, the change-point model becomes less certain about the location of the change-point and the distribution becomes more diffuse. By simplifying dynamics to only mean and amplitude we can capture the main features of the time series dynamics, and additional information as to the speed of the change can be gained from the distributions around the change-points.

Code for this analysis is available on GitHub (https://github.com/emchristensen/Extreme-events-LDA) and archived on Zenodo (Christensen et al. 2018).

## Results

Rodent species composition over the 40 years of the study was best described using four different community-types (Appendix S2: Fig. S1, Table S1). Our four community-types share some species (though they differ in relative abundances), while other species are unique to one or two community-types (Fig. 1a). Our four community-types differ in which species are the most abundant. Community-types 1 and 2 are dominated by kangaroo rats from the genus *Dipodomys*: community-type 1 is co-dominated by *D. spectabilis* and *D. merriami*, while community-type 2 is dominated by *D. merriami* alone. In contrast, the most abundant members of community-types 3 and 4 are pocket mice from the genus *Chaetodipus*: community-type 3 is dominated by *C. baileyi* and community-type 4 by *C. penicillatus*. In Fig. 1a, the 21 species are arranged on the x-axes in order of decreasing body size and grassland-affiliated species are denoted with bold outlines on their bars, to demonstrate that the four community-types differ not only in the identity of species making up the community, but also the distribution of body sizes (Ernest 2013) and habitat preferences contained in the community.

**Figure 1.**
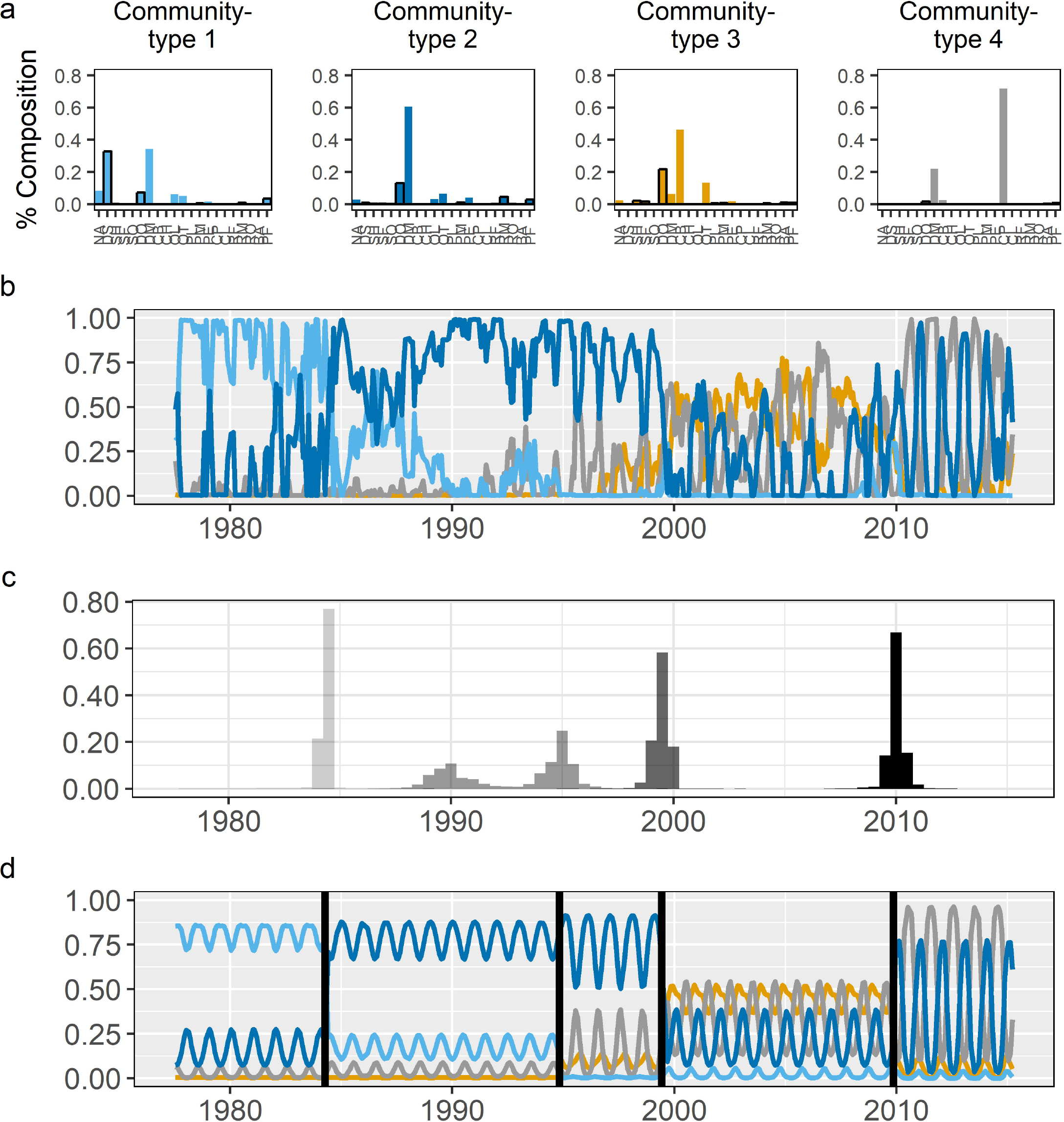
**a)** species composition of the four community-types produced by the LDA model, with species arranged on the x-axes by decreasing body size, and grassland-affiliated species emphasized by black boxes around the bars (see Appendix S2: Table S1); **b)** prevalence of the four community-types over time as estimated by the LDA model; **c)** histograms of four change-points representing the greatest changes in the prevalence of community-types from b; and **d)** the change-point model’s estimate of how community-type prevalence changes before and after each transition point. Species codes in panel a: NA = *Neotoma albigula*, DS = *Dipodomys spectabilis*, SH = *Sigmodon hispidus*, SF = *Sigmodon fulviventer*, SO = *Sigmodon ochrognathus*, DO = *Dipodomys ordii*, DM = *Dipodomys merriami*, CB = *Chaetodipus baileyi*, CH = *Chaetodipus hispidus*, OL = *Onychomys leucogaster*, OT *= Onychomys torridus*, PL = *Peromyscus leucopus*, PM = *Peromyscus maniculatus*, PE = *Peromyscus eremicus*, CP = *Chaetodipus penicillatus*, CI = *Chaetodipus intermedius*, RF = *Reithrodontomys fulvescens*, RM = *Reithrodontomys megalotis*, RO = *Reithrodontomys montanus*, BA = *Baiomys taylori*, PF = *Perognathus flavus*.

Through time, the different community-types varied in their prevalence and dynamics (Fig. 1b). When the study began, the desert rodent community mainly consisted of community-type 1 (Fig. 1, light blue). In the mid-1980s, the rodent community transitioned to community-type 2 (Fig. 1, dark blue) and then transitioned again in the late 1990s to become a mix of community-types 2, 3, and 4 (Fig. 1, dark blue, gold, and grey, respectively). Finally, around 2010, the community entered its current state which is seasonal oscillations between community-types 2 and 4 (Fig. 1, dark blue and grey). These dynamics and community-types are consistent with previous studies that documented the decline of *D. spectabilis* (the co-dominant species of community-type 1) in the mid-1980s (Valone et al. 1995), the colonization and rise to dominance of *C. baileyi* (the dominant species in community-type 3) in the late-1990s (Ernest and Brown 2001), shifts in the body size structure of the community from large species to smaller species (White et al. 2004), and a general decline in grassland-affiliated species and an increase in shrubland-affiliated species (Ernest et al. 2008).

Visually, the LDA results suggest that major shifts in community dynamics occurred multiple times over the study. Using our change-point approach, we found that a model containing four change-points was best supported by the data (for comparison of models containing 2, 3, 4, and 5 change-points see Appendix S2: Fig. S2). Histograms showing the locations of these four change-points are shown in Fig. 2, with the distribution of each point shown in a different shade of gray. Using these distributions, we identified the 95% credible interval for the timing of these transitions: December 1983-July 1984, October 1988-January 1996, September 1998-December 1999, and June 2009-September 2010. Comparison of these distributions with our simulations for different rates of change (Appendix S1: Fig. S2) suggest that three of these events occurred relatively rapidly, i.e. in less than five years. Fig. 1d shows the change-point model’s estimate of how the prevalence of the four community-types differs before and after each transition event, demonstrating that three of these events (1984, 1998-1999, 2009-2010) are driven by a shift in which community-type is most prevalent, marking a major shift in community structure.

**Figure 2.**
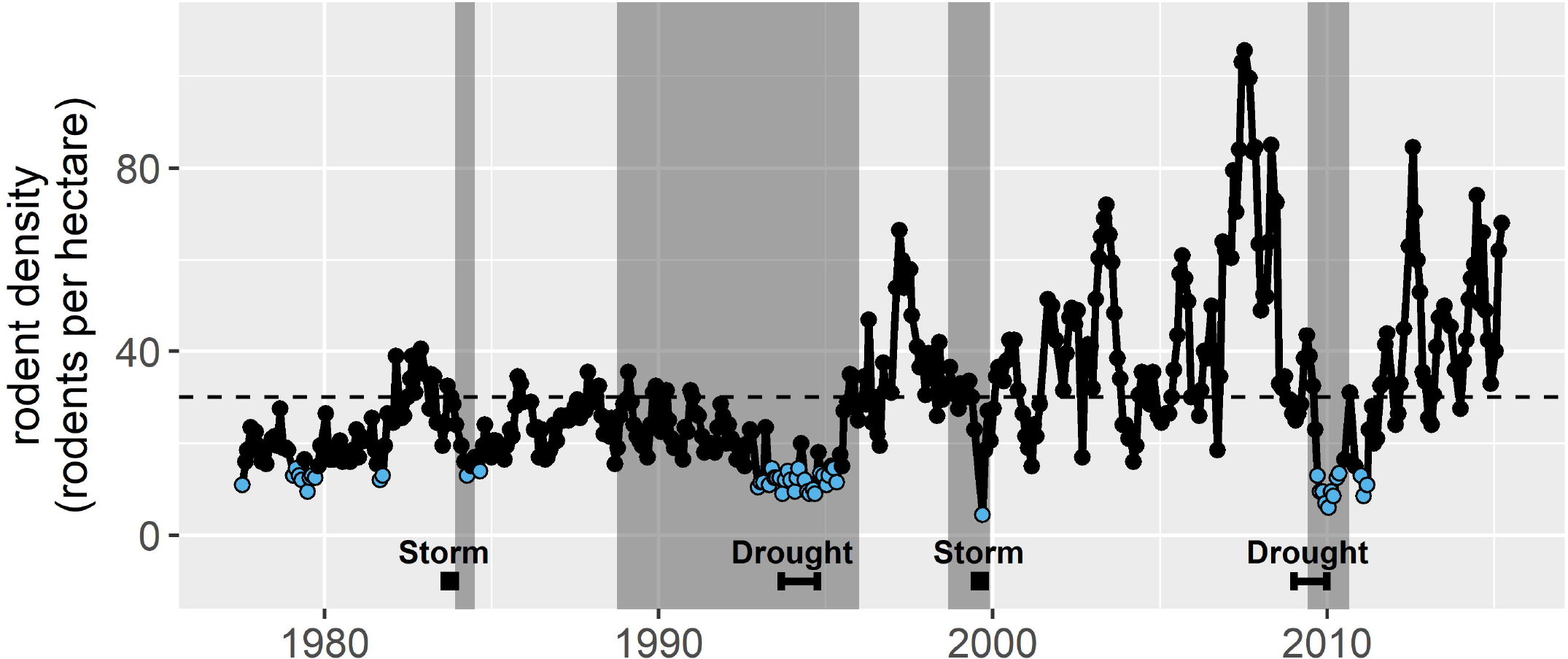
Total rodent abundance per hectare over time. Horizontal dotted line shows long-term mean. Grey vertical bars show the 95% credible interval for each of the community transition events. Light blue dots are data points in the 0.15 probability quantile of the negative binomial distribution fit to the data. Locations in time of the two droughts and two storm disturbance events are shown.

The 1988-1996 transition differs from the other three major reorganization events. It has a broader change-point distribution, and there is no change in which community-type is dominant. However, the change-point model detects an increase in the variance (amplitude) of the dominant community-type through this period and minor changes in community structure as community-type 1 disappears and community-types 3 and 4 increase in prevalence. These dynamics suggest the 1988-1996 transition is not a rapid shift in community structure like the other events, but is instead a period of increased variability in community structure and/or is a signal of a very slow shift in species composition that was abruptly terminated with the major reorganization event of 1998-1999. We also repeated analyses with three, five and six community-types in case our results were highly sensitive to the number of community-types specified, and qualitatively, the results we discuss are generally robust (see Appendix S2: Fig. S3, Fig. S4, Fig. S5 for comparison).

## Discussion

Over nearly 40 years, the rodent community at the Portal Project has changed substantively, with shifts in species composition, dominance structure, and distributions of body sizes and habitat affiliations (Fig. 1). Our results are consistent with earlier studies which described the replacement of grassland-affiliated species by shrubland-affiliated species (Ernest et al. 2008). However, our results indicate that this reorganization did not unfold as a simple gradual event, but a more complicated series of events: a combination of relatively discrete events that occurred roughly every 10-15 years (Fig. 1), and a long period in the 1980s which could indicate a prolonged period of gradual change or increased variance. While the overall shift from a large-bodied grassland-affiliated assemblage to a smaller-bodied shrubland-affiliated assemblage seems to have occurred primarily during concentrated periods, our current analysis does not rule out the possibility of subtle/gradual changes leading up to the periods of more rapid change. The LDA results (Fig. 1b) do seem to show subtle changes in the community leading up to transition events, but quantifying those more subtle dynamics will require development of more sophisticated tools. Altogether, our results show that long-term change in this system was complex, but that rapid shifts were an important and repeated component of long-term change.

Our analysis only quantifies the dynamics of change, not the processes driving them, but we can pull from the natural history of this site and previously published research to provide some hypotheses. The shift in the assemblage from grassland- to shrubland-affiliated species suggests that the documented increase in shrubs (Brown et al. 1997) is likely a significant driver of change, but how it drives rapid transition events is less clear. While we do not have high-frequency observations of this habitat change, shrub growth dynamics are known to be slow compared to the rapid changes we observed in the rodent community (Goslee et al. 2003). When we examine the timing of the rapid transitions, there is a coincidence between the location of the change-point distributions and low abundance periods for the rodents (Fig. 2). Not all low abundance events are associated with rapid shifts in composition, but all rapid shifts in composition are associated with low abundance events. The abundance of an entire community can drop for a variety of reasons (e.g., low resource availability, disturbances, disease, or predation events) and we do not have the data to examine all possibilities. However, as in many ecosystems, our site experiences both periodic droughts and extreme rainfall events. Droughts reduce resource availability in this water-limited system, and extreme rainfall events can cause sheet flooding (Thibault and Brown 2008) or saturate soils and damage food stores for granivorous rodents (Valone et al. 1995). All four of our detected change-points, including the longer transition in the 1990s, overlap or occur adjacent to droughts or high rainfall events: 1) an intense tropical storm in October 1983 (Valone et al. 1995), 2) a drought in the 1990s (Allington et al. 2013; Appendix S2: Fig. S6), 3) a sheet flood during monsoon season in August 1999 (Thibault and Brown 2008), and 4) a period of low plant productivity in 2009 (Appendix S2: Fig. S6). However, it is important to note that not all droughts and high rainfall events in this system are associated with low abundance events. For example, low plant productivity also occurred in 2003 (Appendix S2: Fig. S6), but neither abnormally low abundances nor a transition in community composition occurred during that time. While correlation does not prove causation, the co-occurrence of low abundances and transition periods with climate events suggests that studying these linkages further may be productive.

There are a variety of processes that could link low abundance to an increased likelihood of a rapid ecological transition. One hypothesis for why low abundance could generate transition events is that low abundances occur when the dominant species has been hit disproportionately hard, releasing other species from competition and opening niche opportunities (Shea and Chesson 2002). This is a possible explanation for our 1983 transition event when the dominant species *D. spectabilis* experienced a population crash (Valone et al. 1995). The loss of this species may also explain the extended 1988-1996 transition period: *D. spectabilis* is also an ecosystem engineer, constructing large burrow mounds which are used by a variety of animals and altering the plant community (Guo 1996), and the loss of this species may have had indirect and/or lagged effects on the rodent community. Alternatively, low community abundance can generate new community compositions by increasing the role of stochasticity in community assembly through either its impacts on competitive outcomes (Orrock and Fletcher 2005), or by creating stochastic opportunities for colonization (Fukami et al. 2010). Events such as the 1999 flood, when all members of the community decrease to similarly low numbers (Thibault and Brown 2008), may be critical junctures where a variety of stochastic forces may operate to drive communities toward new assembly trajectories. However, given that the niche characteristics of our species appear to be tracking the shift in environment, it seems unlikely that stochastic processes alone explain why our community structure changes after low abundance events. A third hypothesis is that priority effects slow the ability of a community to track environmental change (like slowly increasing shrub cover) and that low abundance events allow the community to rapidly “catch up” to the environment by removing these priority effects. The ability of an established community to resist new colonists (Amarasekare 2002) is well-documented in the context of invasive species (Corbin and D’Antonio 2004) but its role in determining how communities track environmental change has received less attention (but see Thibault and Brown 2008). As our environment shifted from grassland to shrubland, species that were competitively dominant would have slowly become competitively inferior to shrubland species, but the “incumbent advantage” may have impeded the ability of superior competitors to colonize and increase (Amarsekare 2002). Reductions in abundance would remove this “incumbent advantage” by creating a clean slate where superior competitors can now dominate a community. At this time, it is not possible to disentangle if any, all, or none of these processes are behind our patterns of change, but further exploration of the link between low abundance events and rapid transitions seems worthwhile given the number of pathways through which low abundance could trigger rapid change.

While we cannot determine what processes are behind the patterns of change in our rodent community, our results show that the pattern of long-term community change was not simple. In this system, much of the long-term change in composition occurs in concentrated bursts, followed by periods of little, or at least much slower, change. Given our methods, we are constrained to make relative and qualitative statements about how quickly “rapid” change occurred, but change is a continuous rate that should be measurable. Clearly, a major goal for future methods development is not just documenting qualitative patterns of change, but also quantifying rates of change and how those rates differ through time. While better methods are being developed, our research shows that beginning to understand the patterns of long-term change is an important immediate step. Whether long-term change commonly occurs through multiple rapid events or if this pattern is unique to our system is critical for understanding whether systems that appear stable today may be on the verge of rapid ecological transitions. However, rectifying long-term multidecadal scale changes with the short-term dynamics that create that change requires long-term, high frequency monitoring, emphasizing growing concerns (Hughes et al. 2017) that maintaining long-term studies will be critical for detecting, understanding, and predicting future changes in nature.

## Acknowledgements

The Portal Project has been funded nearly continuously by the National Science Foundation, most recently by DEB-1622425, which also supported E. Christensen. This research was also supported in part by the Gordon and Betty Moore Foundation’s Data-Driven Discovery Initiative through Grant GBMF4563 to E.P. White. We would also like to thank E. Damschen for informative discussions on stochastic community assembly, and D. Valle and J. Simonis for insights on the LDA method.

